# Novel, non-nitrocatechol COMT inhibitors modulate dopamine neurotransmission in the frontal cortex and improve cognitive flexibility

**DOI:** 10.1101/510040

**Authors:** Spencer Byers, Ingrid P. Buchler, Michael DePasquale, Helen L. Rowley, Rajiv S. Kulkarni, Lucy Pinder, Anna Kolobova, Cailian Li, Vinh Au, Daniel Akuma, Gongliang Zhang, Huijun Wei, Sharon C. Cheetham, James C. Barrow, Gregory V. Carr

## Abstract

Cognitive impairment is a primary feature of many neuropsychiatric disorders and there is a need for new therapeutic options. Catechol-*O*-methyltransferase (COMT) inhibitors modulate cortical dopaminergic function and have been proposed as potential cognitive enhancers. Unfortunately, currently available COMT inhibitors are not good candidates due to either poor blood-brain barrier penetration or severe toxicity. To address the need for safe, brain-penetrant COMT inhibitors, we tested multiple novel COMT inhibitors in a set of preclinical *in vivo* efficacy assays to determine their viability as potential clinical candidates. We found that multiple COMT inhibitors, exemplified by LIBD-1 and LIBD-3, significantly modulated dopaminergic function measured as decreases in homovanillic acid (HVA) and increases in 3,4-Dihydroxyphenylacetic acid (DOPAC), two dopamine metabolites, in cerebrospinal fluid (CSF) and the frontal cortex. Additionally, we found the LIBD-1 significantly improved cognitive flexibility in a rat attentional set-shifting assay (ASST), an effect previously seen with the COMT inhibitor tolcapone. These results demonstrate that LIBD-1 is a novel COMT inhibitor with promising *in vivo* activity and the potential to serve as a new therapy for cognitive impairment.

## Introduction

Cognitive impairment is a core feature of many neurological and psychiatric disorders and the severity of the impairment is directly correlated with functional outcomes for patients (1-5). Currently, there are very few therapeutic options for patients with cognitive impairment. The options that are available have limited efficacy and in some cases, including psychostimulants in schizophrenia, are contraindicated due to potential exacerbation of other symptoms (6).

The neuromodulator dopamine (DA) plays a critical role in cognitive function across species and strategies that augment or refine DA activity in the brain have been shown to improve cognitive function (7). The enzyme catechol-*O*-methyltransferase (COMT) is a key component of the DA metabolic pathway (8) and inhibition of the enzyme has been proposed as a cognitive enhancement strategy (9). COMT is located throughout the body with the highest expression in the liver and brain (10). There is a soluble (S-COMT) and a membrane-bound (MB-COMT) form of the enzyme. S-COMT has a higher Vmax but lower affinity for DA compared to MB-COMT (11). It is thought that S-COMT activity predominates in the periphery where it metabolizes many substrates including catecholestrogens and exogenous catechols (12). In contrast, MB-COMT accounts for most of the enzyme activity in the brain, especially in the primate brain, where the major substrate is dopamine, which is present at relatively low concentrations (13).

The major routes for terminating DA signaling in the brain are reuptake and metabolic conversion. In the striatum, the dopamine transporter (DAT) is the major termination mechanism. In the frontal cortex and hippocampus, there are relatively small amounts of DAT, most of which is extrasynaptic. There is evidence that the norepinephrine transporter (NET) can clear DA from the extracellular space in cortex and hippocampus (14), but NETs also are outside DA synapses and the majority of functional DA signal termination in these regions appears to depend on COMT activity, with some contribution from monoamine oxidase (15, 16). COMT directly converts DA to 3-Methoxytyramine (3-MT) and also converts the DA metabolite 3,4-Dihydroxyphenylacetic acid (DOPAC) to homovanillic acid (HVA). The fact that COMT has an outsized role in DA signaling in the cortex and hippocampus compared to the striatum has led to the hypothesis that selective targeting of COMT activity may modulate cognitive function without some of the psychostimulant effects (e.g. abuse liability) seen with other DA modulators (9).

There is a large body of evidence showing that COMT activity has a dynamic relationship with cognitive function in humans and animal models. There is a highly-studied functional polymorphism in the COMT gene (Val158Met) that produces a “high” activity enzyme (Val158) and a “low” activity enzyme (Met158). In general, the Val form of COMT is associated with lower cognitive function compared to the Met form (17-19). However, this effect is labile and interactions with other genetic and environmental factors can modify or even reverse the relationship (20, 21). Administration of COMT inhibitors also has variable effects on cognitive function based on the individual status of the Val/Met polymorphism and baseline performance, particularly in studies of people without a diagnosis of cognitive impairment (22, 23). Overall, there have been a number of studies of patients across multiple indications that suggest that COMT inhibition may be a viable strategy for improving cognitive function (24, 25).

Currently, COMT inhibitors are utilized as an adjunctive therapy alongside L-DOPA administration in patients with Parkinson’s disease (PD) to improve motor symptoms. COMT metabolizes L-DOPA to 3-*O*-Methyldopa (3-OMD) in the periphery and decreases the amount of L-DOPA available to cross the blood-brain barrier (BBB). Therefore, COMT inhibition serves to augment the CNS availability and effects of L-DOPA without having to increase the dosage of L-DOPA (26, 27). Since the target enzyme is in the periphery, a brain-penetrant COMT inhibitor is not necessary for this indication. However, for cognitive enhancing effects, there is evidence that a brain-penetrant or central COMT inhibitor is required. A comparison of the effects of tolcopone (brain-penetrant) to those of entacapone (peripherally-restricted) in patients with PD showed that only those taking tolcapone demonstrated improvement in the cognitive symptoms associated with PD (24).

Tolcapone, however, is not a viable, long-term option as a cognitive enhancer due to safety concerns. Tolcapone is associated with potentially fatal liver toxicity that makes the drug a poor choice for a chronic cognitive enhancement therapy (28). Given the relative safety of the other COMT inhibitors on the market, the toxicity is thought to be compound-specific as opposed to mechanism-based (29). Therefore, a potent, brain-penetrant, and safe COMT inhibitor is needed to fully explore this mechanism as a potential avenue for cognitive enhancement.

Here we describe the *in vivo* activity of novel COMT inhibitors. Our goal was to identify a COMT inhibitor with good oral bioavailability and significant effects on neurochemical biomarkers and behavior in rats. We used tolcapone as our efficacy benchmark. We identified LIBD-1 as a candidate molecule for future studies. LIBD-1 has high selectivity for MB-COMT over S-COMT, which we believe will produce an outsized effect on central COMT activity while having a smaller impact on liver COMT function. The evidence presented here indicates that LIBD-1 is a viable lead compound with favorable properties that support continued development.

## Materials and Methods

### Animals

Male CD^®^ IGS rats (8-11 weeks old; Charles River Laboratories, Wilmington, MA, USA) were used for all experiments. Age-matched female CD^®^ IGS rats were also used for some of the DA metabolite biomarker studies. Rats used for experiments at the Lieber Institute for Brain Development were housed in a temperature (22°C) and humidity-controlled facility on a 12h/12h light/dark cycle (lights on at 0600 hours). Rats used for experiments at RenaSci were housed in a temperature (21 ± 4°C) and humidity-controlled (55 ± 15%) room on a 12h/12h light/dark cycle (lights on at 0700 hours) facility. Rats used in behavioral experiments were handled for one minute at least three times in the week prior to testing. All procedures were approved by the local Animal Care and Use Committee and were in compliance with the *Guide for the Care and Use of Laboratory Animals*.

### Drugs

Novel COMT inhibitors (synthesized in-house) and tolcapone (synthesized in-house) were suspended in vehicle (0.1% Tween80, 0.1% 1510 silicone antifoam, 1% methylcellulose in water; Table 1) and administered either orally (po) or through intraperitoneal (ip) injection (tolcapone). Administration volumes were 10 mL/kg and 5 mL/kg for po and ip dosing, respectively. In the microdialysis experiment, a dose volume of 2 mL/kg was used for tolcapone administration.

| Compound | Human MB-COMT IC50 (nM) | Human S-COMT IC50 (nM) | Rat MB-COMT IC50 (nM) | Rat S-COMT IC50 (nM) |
| --- | --- | --- | --- | --- |
| Tolcapone | < 1 | < 1 | < 1 | < 1 |
| LIBD-1 | 5.8 | 1906 | < 1 | 3750 |
| LIBD-2 | 7.7 | 3.8 | < 1 | 236 |
| LIBD-3 | 1.5 | > 5000 | 2.2 | > 5000 |

All COMT potency values are for recombinant enzyme
LIBD-1 and LIBD-2 are 8-Hydroxyquinolines from the family of compounds described in reference 32
LIBD-3 is a pyrido[1,2-*a*]pyrazine-8-one described in patent WO2017091818

### DA metabolite biomarker protocol

On the day of testing, rats were transferred to a holding room and weighed. After an hour acclimation period, rats received either vehicle or a COMT inhibitor. At the appropriate time point (1 or 4 hours), rats were moved to a separate procedure room where they were anaesthetized via isoflurane. Once the rats were determined to be unresponsive, their heads were shaved using electric clippers. The rats were positioned in a stereotaxic frame, with their heads pointed down at a 45-degree angle. To collect cerebrospinal fluid (CSF), previously published protocols (30, 31) were adapted. Briefly, a 23 gauge needle, connected via PE50 tubing to a collection syringe, was used to access the cisterna magna. Slight negative pressure was used to ensure the CSF flowed evenly. The CSF was collected in previously chilled (dry ice) Eppendorf tubes containing 0.05M perchloric acid (4:1 CSF:perchloric acid ratio). The tubes were put back on dry ice until the end of the procedure. CSF samples with visible blood contamination were not used in subsequent bioanalytical analyses. Next, the chest cavity was opened, and blood was collected by cardiac puncture. The blood was collected in Lithium-Heparin 1.3mL microtubes (Sarstadt, Numbrecht, Germany) and stored on ice. The rat was then transcardially perfused with ice-cold phosphate buffered saline (PBS) via a peristaltic pump set to a flow rate of 20 mL/min. Following perfusion, the brains were collected in 5mL Thermo Scientific Nunc tubes (Thermo Fisher Scientific, Waltham, MA, USA). Blood was then centrifuged at 2000 g at 4°C for 10 minutes to separate the plasma. Plasma was then transferred into Thermo Scientific Matrix tubes (Thermo Fisher Scientific, Waltham, MA, USA) for storage. CSF, plasma, and brains were stored at −80°C until analysis. DA metabolite and COMT inhibitor concentrations were measured by LC-MS/MS as previously described (32).

### Frontal cortex microdialysis

#### Surgery

Rats were anaesthetised with isoflurane (5% to induce, 2% to maintain) in O2 (1 L/min) delivered via an anaesthetic unit (Burtons Medical Equipment Ltd, UK). A concentric microdialysis probe (CMA 12 Elite, with an exposed polyarylethersulphone (PAES) membrane tip; Linton Instrumentation, Norfolk, UK) was stereotaxically implanted into the frontal cortex (2 mm tip, coordinates: AP: +3.2 mm; L: +/-2.5 mm relative to bregma; V: −4.0 mm relative to the skull surface). Coordinates were taken from the stereotaxic atlas of Paxinos and Watson (1986). The upper incisor bar was set at 3.3 mm below the interaural line so that the skull surface between bregma and lambda was horizontal. Additional burr holes were made for skull screws (stainless steel) and the probes were secured using dental cement. Carprofen (Carprieve, Norbrook Laboratories (GB) Ltd, Newry, UK) was administered for pain relief at least 30 min prior to animals regaining consciousness following surgery (5 mg/kg sc). Following surgery, animals were individually housed in the dialysis bowls (245 mm internal diameter at base of bowl, 360 mm wall height, Harvard Apparatus, Cambridge, UK), with the microdialysis probes connected to swivels and a counter-balanced arm to allow unrestricted movement. Rats were allowed a recovery period of at least 16 h with standard rodent chow and filtered water available ad libitum. During this time, the probes were continuously perfused at a flow rate of 1.2 μL/min with an artificial cerebrospinal fluid (aCSF; Harvard Apparatus, UK) of the following electrolyte composition (in mM): sodium 150; potassium 3.0; magnesium 0.8; calcium 1.4; phosphate 1.0; chloride 155.0, pH 7.2.

### Dialysate sample collection

Dialysate samples were collected at 20 min intervals for a baseline period of 80 min prior to the onset of administration of vehicle, tolcapone, LIBD-1, or LIBD-3. Sampling continued for an additional 2 h. Samples were collected into Eppendorf vials (300 μl, Microbiotech, Stockholm, Sweden) containing 5.0 μL of 0.1 M perchloric acid to prevent oxidation of DA and its metabolites. Samples were stored at −80°C until assayed by HPLC with electrochemical detection.

Additional details on bioanalytical analysis and microdialysis probe location verification are provided in Supplemental Information.

### Attentional set-shifting task (ASST)

This version of the ASST was adapted from previously described protocols (33, 34), and was performed over a four-day period. Rats were food restricted to 90% of free-feeding weight during the week before testing. A full description of the methods is provided in the Supplemental Material.

### Data analyses

In our CSF neurochemical experiments, we compared drug-treated rats to vehicle-rats using either one-way ANOVAs or t-tests. For ANOVAs significant main effects were followed up with post hoc Tukey’s tests. Data are represented by the mean value as a percentage of the vehicle-treated group. For the microdialysis experiment, data were log transformed and analyzed by ANCOVA with log(baseline) as a covariate. Baseline was defined as the geometric mean of the four pretreatment samples. Comparisons to the vehicle-treated group were by the multiple t-test (Supplemental Tables 2-4). Data are presented as the mean (% of vehicle-treated) ± SEM. Some extreme values were removed from the analysis. These are listed in Supplemental Table 5. The ASST data were analyzed using two-way ANOVA with repeated measures. Drug treatment was the between-subjects variable and ASST stage was the within-subjects variable. Significant main effects were further probed with post hoc Tukey’s tests to determine significant pairwise comparisons. ASST data are presented as the mean ± SEM.

## Results

### COMT inhibitors modulate HVA and DOPAC concentrations in CSF

To determine if our novel COMT inhibitors (Table 1) prevent COMT activity *in vivo*, we measured the concentrations of the dopamine metabolites HVA and DOPAC in cerebrospinal fluid (CSF). Previous research has shown that potent brain-penetrant COMT inhibitors reliably decrease HVA concentrations while increasing DOPAC concentrations in CSF (35). Here, we benchmarked our COMT inhibitors to the prototypical COMT inhibitor tolcapone (Table 2). Tolcapone (15 mg/kg) significantly decreased HVA (1 h: *t_9_* = 8.730, *p* < 0.0001; 4 h: *t_13_* = 13.090, *p* < 0.0001) and increased DOPAC (1 h: *t_9_* = 8.528, *p* < 0.0001; 4 h: *t_13_* = 9.866, *p* < 0.0001) at both time points sampled (1 and 4 h). All doses of our MB-COMT selective inhibitors LIBD-1 and LIBD-3 also significantly decreased HVA (LIBD-1 1 h: *F*_3,30_ = 26.450, *p* < 0.0001; 4 h: *F*_3,33_ = 29.470, *p* < 0.0001; LIBD-3 1 h: *F*_3,32_ = 42.700, *p* < 0.0001; 4 h: *F*_2,19_ = 126.200, *p* < 0.0001) and increased DOPAC (LIBD-1 1 h: *F*_3,30_ = 46.850, *p* < 0.0001; 4 h: *F*_3,33_ = 42.670, *p* < 0.0001; LIBD-3 1 h: *F*_3,32_ = 32.730, *p* < 0.0001; 4 h: *F*_2,19_ = 41.510, *p* < 0.0001) at both time points tested. The highest doses of LIBD-1 and LIBD-3 were unable to modulate the DA metabolites to the same degree as tolcapone despite reaching concentrations that would be expected to completely block MB-COMT, suggesting some role for S-COMT in regulating DA metabolism in rats. Previous research has shown sexually dimorphic effects related to COMT activity (36, 37), so we tested both tolcapone (HVA: *t_10_* = 9.505, *p* < 0.0001; DOPAC: *t_10_* = 6.307, *p* < 0.0001) and LIBD-1 (HVA: *t_11_* = 6.696, *p* < 0.0001; DOPAC: *t_11_* = 5.478, *p* = 0.0003) in female rats and found similar changes in DA metabolites. In contrast, LIBD-2, a potent, isoform nonselective COMT inhibitor, with extremely low brain penetrance, does not significantly alter CSF DA metabolites (HVA: *t_13_* = 0.1482, *p* = 0.8845; DOPAC: *t_13_* = 1.793, *p* = 0.0962), indicating that our biomarker assay is a sensitive measure of *in vivo* target engagement.

| Compound | Dose | Timepoint (h) | Free Plasma (nM) | Free Brain (nM) | CSF (nM) | CSF HVA (%vehicle) | CSF DOPAC (%vehicle) |
| --- | --- | --- | --- | --- | --- | --- | --- |
| Tolcapone | 15 mg/kg | 1 | 35 | BLQ | BLQ | 7 | 291 |
|  | 15 mg/kg | 4 | 4 | BLQ | BLQ | 28 | 304 |
|  | 15 mg/kg (IP); females | 4 | 10 | BLQ | BLQ | 31 | 233 |
| LIBD-1 | 10 mg/kg | 1 | 69 | 62 | 132 | 63 | 234 |
|  | 30 mg/kg | 1 | 98 | BLQ | 352 | 49 | 229 |
|  | 100 mg/kg | 1 | 211 | 191 | 1220 | 36 | 273 |
|  | 10 mg/kg | 4 | 35 | 35 | 157 | 72 | 140 |
|  | 30 mg/kg | 4 | 64 | BLQ | 176 | 66 | 210 |
|  | 100 mg/kg | 4 | 175 | 67 | 423 | 49 | 251 |
|  | 100 mg/kg; females | 4 | 142 | 128 | 349 | 42 | 238 |
| LIBD-2 | 100 mg/kg | 4 | 41 | BLQ | BLQ | 101 | 119 |
| LIBD-3 | 10 mg/kg | 1 | N/A | N/A | N/A | 59 | 201 |
|  | 30 mg/kg | 1 | 145 | 156 | 84 | 39 | 197 |
|  | 100 mg/kg | 1 | 496 | 674 | 328 | 35 | 276 |
|  | 30 mg/kg | 4 | 46 | 39 | 15 | 48 | 182 |
|  | 100 mg/kg | 4 | 290 | 430 | 78 | 22 | 207 |

BLQ = below limit of quantitation
Limits of quantitation: Tolcapone (74 nM in CSF and 144 nM in brain). LIBD-1 (66 nM in brain and CSF). LIBD-2 (58 nM in CSF and 233 nM in brain).
N/A = no values

### COMT inhibitors modulate HVA and DOPAC concentrations in the frontal cortex

The CSF biomarker assay is a good measure of target engagement, but we also wanted to confirm the activity of our compounds in the putative brain subregion, the cerebral cortex, thought to be the neural substrate for the effects of COMT inhibition on cognitive function. Therefore, we also tested the effects of our novel COMT inhibitors on DA, HVA, and DOPAC concentrations in extracellular fluid (ECF) sampled from the cortex by microdialysis (Figure 1). As previously reported, none of the COMT inhibitors altered extracellular DA concentrations (38-40). Here, both LIBD-1 and LIBD-3 significantly decrease HVA and increase DOPAC as expected (Figure 1b-c). As seen in the CSF experiment, LIBD-1 effects do not quite reach the same magnitude as our positive control tolcapone. In contrast, the effects of LIBD-3 were not significantly different from tolcapone. These microdialysis results support the effects we see in CSF and further demonstate that our MB-selective compounds may have similar utility as compared to tolcapone.

**Figure 1.**
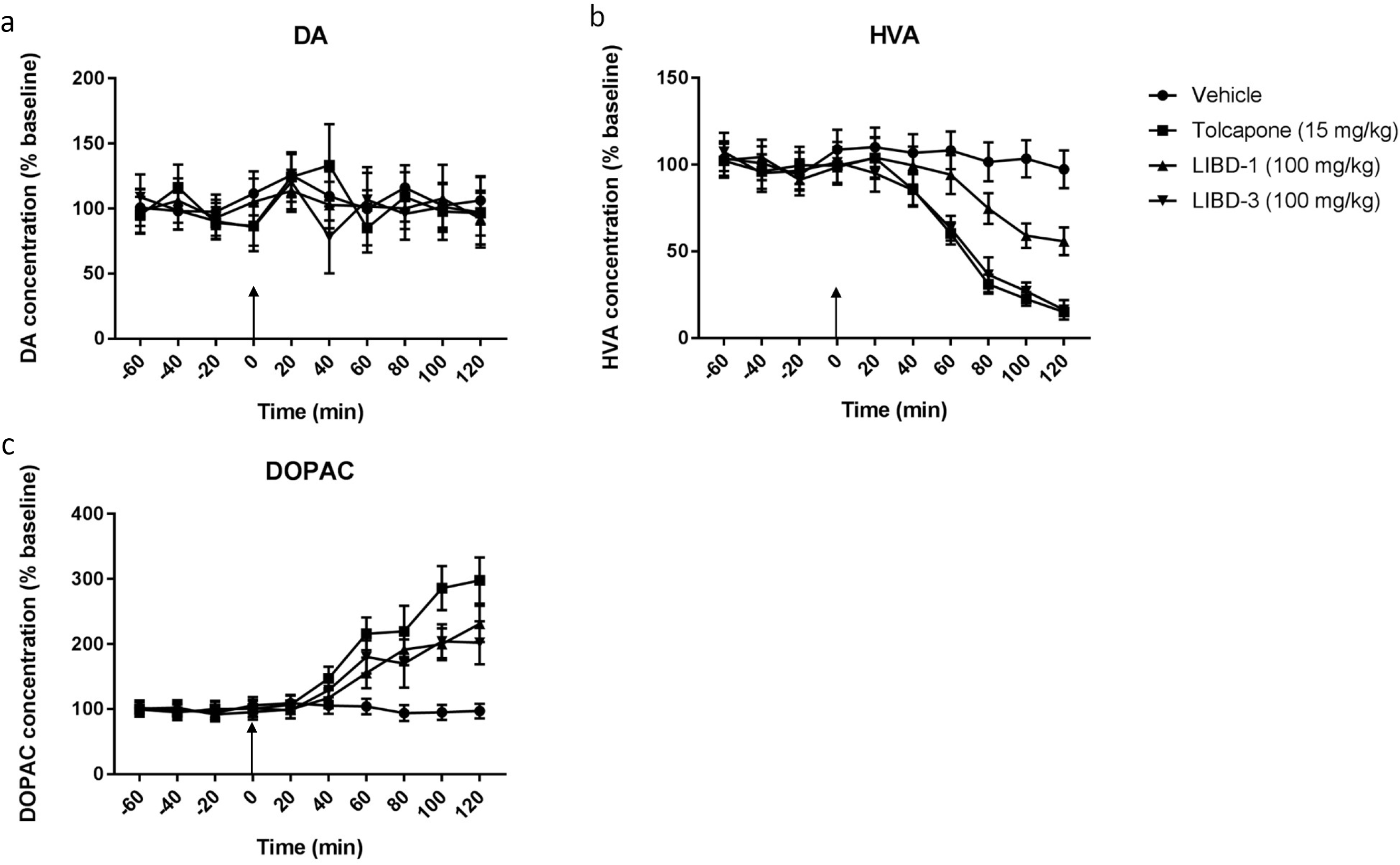
Effects of COMT inhibitors on DA and metabolites in the frontal cortex. **a** DA concentration. **b** HVA concentration. **c** DOPAC concentration. Results are adjusted means; n=8 for all groups except tolcapone (n = 7). Vertical arrow indicates time of drug administration (t = 0). Tolcapone, LIBD-1, and LIBD-3 all decreased HVA concentrations and increased DOPAC concentrations without significantly altering DA concentrations compared to vehicle treatment. Statistical results, including timepoints with significant differences between groups, are presented in Supplemental Tables 1-3.

### LIBD-1 improves cognitive flexibility in the set shifting assay

Both LIBD-1 and LIBD-3 demonstrate good *in vivo* COMT inhibitor activity. The next step was to test whether LIBD-1 and LIBD-3 would be able to modulate cognitive function, measured as performance in the ASST. The rat digging version of the ASST is a validated assay for measuring cognitive flexibility and the task is dependent on an intact frontal cortex (33). Moreover, COMT inhibition has been shown to improve performance during the extradimensional shift (ED) stage of the task (38), analogous to the task in monkeys that has been shown to be a dopamine mediated prefrontal cognitive task (41). To establish the validity of our in-house protocol, we first measured the effects of tolcapone (15 mg/kg) as a positive control. As expected, there were significant differences in the number of trials to criterion between the stages (*F*_6,126_ = 4.729, *p* = 0.0002) and tolcapone improved performance (*F*_1,2_ = 5.075, *p* = 0.0351); Figure 2a). Post hoc analyses revealed that the ED stage required more trials to criterion than the compound discrimination stage (CD; *p* = 0.0104) and the intradimensional shift stage (ID; *p* = 0.0003) in vehicle-treated rats. Tolcapone improved ED stage performance (*p* = 0.0177) without significantly altering performance on any of the other stages.

**Figure 2.**
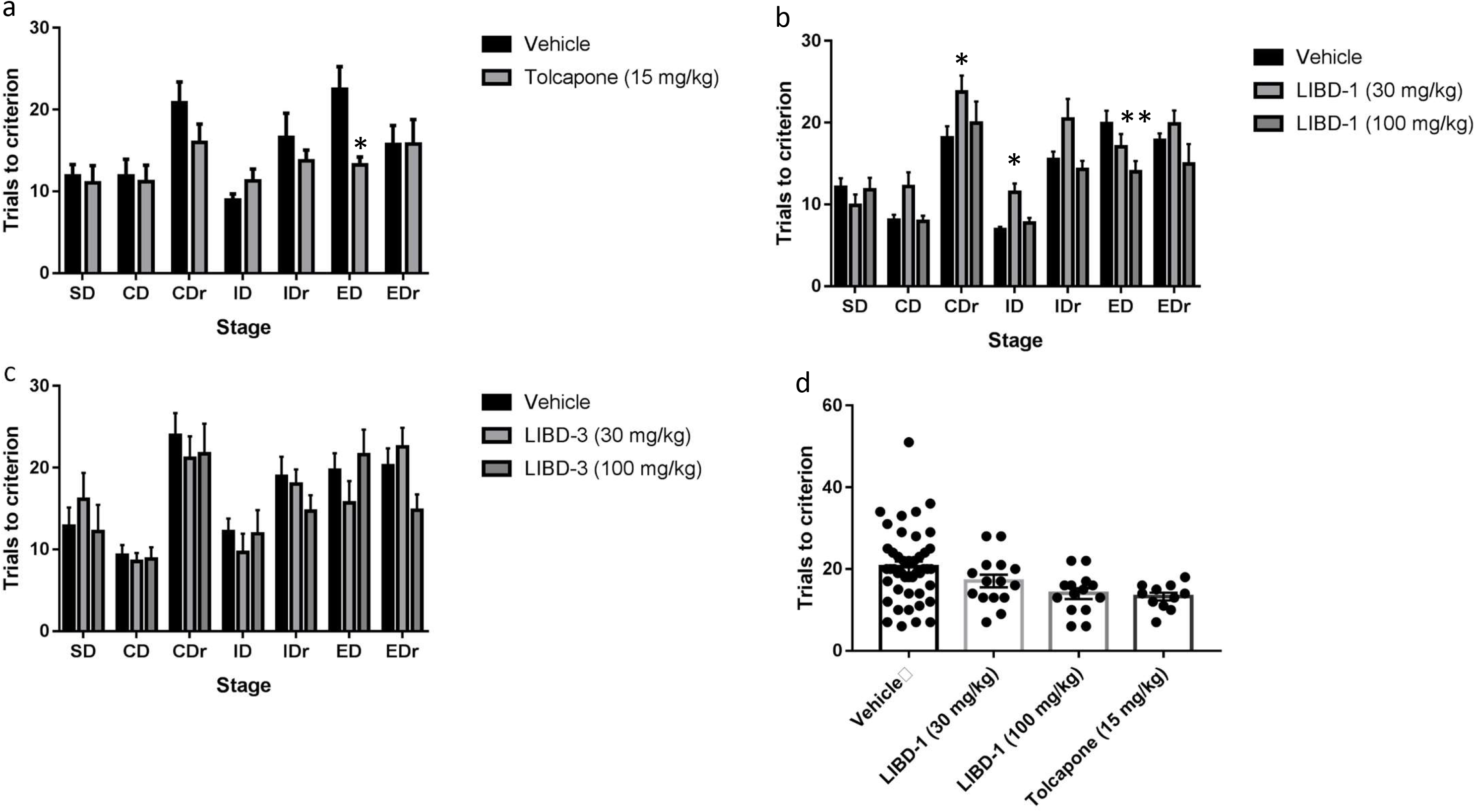
Effects of COMT inhibitors on behavior in the ASST. **a** Tolcapone (n = 12/vehicle and n = 11/tolcapone **b** LIBD-1 (n = 24/vehicle, n = 15/LIBD-1 30 mg/kg group, and n = 14/LIBD-1 100 mg/kg group). **c** LIBD-3 (n = 21/vehicle, n = 14/LIBD-3 30 mg/kg group, and n = 10/ LIBD-3 100 mg/kg group). **d** Comparison of performance in the ED stage across the three experiments. \**p* <0.05 and \*\**p* <0.01 compared to the vehicle-treated group in the stage.

Next, we tested the effects of LIBD-1 (30 and 100 mg/kg) in the ASST. Again, there was a significant difference in the number of trials to criterion between stages (*F*_6,300_ = 31.8, *p* < 0.0001). In the vehicle-treated rats, there was a significant difference in trials to criterion between the ED stage and the CD (*p* < 0.0001) and ID (*p* < 0.0001) stages. Like tolcapone, LIBD-1 improved performance in the ASST compared to vehicle (*F*_2,50_ = 7.52, *p* = 0.0014; Figure 2b). LIBD-1 (100 mg/kg) significantly improved performance in the ED stage of the task (*p* = 0.0087). Additionally, the 30 mg/kg group performed worse than the vehicle group on the compound discrimination reversal (CDr; *p* = 0.0123) and worse than both the vehicle and 100 mg/kg groups on the intradimensional shift reversal stage (IDr; *p* = 0.0296 and *p* = 0.0142, respectively). These data indicate that LIBD-1 modulates cognitive function and the 100 mg/kg dose produces effects similar to the FDA-approved COMT inhibitor tolcapone.

Finally, we tested the effects of LIBD-3 in the ASST (Figure 2c). There was a significant difference in trials to criterion between the stages (*F*_6,252_ = 12.83, *p* = 0.0001). The ED stage required more trials than the CD and ID stages, although only the differences with the CD stage reached the significance threshold (*p* = 0.0019 and *p* = 0. 0674. respectively). Unlike both tolcapone and LIBD-1, LIBD-3 did not affect performance (*F*_2,42_ = 0.4392, *p* = 0.6475).

## Discussion

Research suggests that COMT inhibition is a viable strategy for improving executive function, with potential utility across many clinical indications (42, 43). Data from patients with Parkinson’s disease indicates that the cognitive benefits of COMT inhibition require a brain-penetrant compound (24). Of the FDA-approved COMT inhibitors, tolcapone has marginal brain penetration. Unfortunately, long-term use of tolcapone is associated with potentially fatal liver toxicity in some patients, therefore, novel, safe, and brain-penetrant COMT inhibitors are needed to explore the utility of the mechanism for cognitive enhancement. Here we describe the in vivo activity of novel, non-nitrocatechol, brain-penetrant COMT inhibitors in rats and identify our current lead drug candidate.

In the CNS, COMT inhibitors are known to alter the concentrations of the DA metabolites HVA and DOPAC. Specifically, COMT inhibition decreases HVA and increases DOPAC concentrations. Previous research has used post-administration measurement of CSF HVA and DOPAC as *in vivo* markers of COMT inhibition (35). We found that two of our novel compounds, LIBD-1 and LIBD-3, produced sustained modulation of HVA and DOPAC concentrations in CSF. Additionally, we showed that the changes in CSF HVA and DOPAC were due to central COMT inhibition by testing the effects of a third compound, LIBD-2. LIBD-2 is extremely potent *in vitro*, but its brain-penetrance is negligible and it did not significantly alter central DA metabolism.

While CSF DA metabolites are translational biomarkers for central COMT inhibition, we believe the ultimate target for the use of COMT inhibitors as cognitive enhancers is the frontal cortex. To determine if our novel COMT inhibitors could modulate COMT function directly within our proposed site of action, we conducted microdialysis experiments with probes sampling from the frontal cortex. We found that both LIBD-1 and LIBD-3 significantly altered HVA and DOPAC concentrations within the cortex similar to the effect produced by tolcapone. Our results confirmed previously reported findings for tolcapone and show that non-nitrocatechol COMT inhibitors produce relevant changes in DA metabolism in the cortex.

The experiments discussed so far showed that our novel COMT inhibitors produce qualitatively similar effects compared to the COMT inhibitor tolcapone. We also confirmed that central measures of COMT inhibition require a brain-penetrant inhibitor. The final characteristic we wanted in a potential lead compound was the ability to modulate behavior in a cognition assay. In the next series of experiments, we showed that LIBD-1 produced behavioral changes at relevant central exposure levels. LIBD-1 improves cognitive flexibility in the ED phase of the task comparable to the effects of tolcapone. Interestingly, the lower dose of LIBD-1 increased the number of trials required to complete the CDr and ID stages. Although it is not clear what caused this change in performance, one hypothesis generated from the clinical literature suggests that tolcapone can impair performance by increasing cognitive control at times when more automatic responding is advantageous (22). This may be a particular problem for “normal” subjects, like the rats in our experiments, compared to those with cognitive impairment. Taken together, these results indicate that our novel non-nitrocatechol COMT inhibitor possesses many of the *in vivo* properties required to test its utility as a cognitive enhancer in a clinical trial.

LIBD-3 produced similar, and arguably superior, results in the neurochemical assays compared to LIBD-1, but did not significantly improve cognitive flexibility in the ASST. At this time, it is not clear what is responsible for the discrepancy in performance. In a cell-based assay, LIBD-3 produces significant cellular toxicity and maximum changes in DOPAC concentrations much lower than LIBD-1 and tolcapone (G. Zhang, personal communication). These data suggest that there may be compound-specific effects that could impede performance in the ASST, but are not evident given the limited temporal and spatial resolution of our *in vivo* neurochemical measures.

A large body of evidence suggests that COMT inhibition improves cognitive function across multiple clinical domains (24, 25, 44). Although there are multiple FDA-approved COMT inhibitors, none of them are suitable for long-term use as cognitive enhancers. Entacapone and opicapone (approved in Europe) do not cross the blood-brain barrier. Tolcapone, which is brain-penetrant, has a black-box warning due to potentially fatal liver toxicity. The toxicity associated with tolcapone appears to be compound-specific, as no other marketed COMT inhibitor shares this liability (29). The lack of an approved, brain-penetrant, and safe COMT inhibitor has prevented the full exploration of the utility of the mechanism for cognitive disorders.

In contrast with tolcapone, which is a nonselective inhibitor, our novel COMT inhibitors are selective for MB-COMT compared with S-COMT. In the rat brain, MB-COMT and S-COMT protein levels are approximately equal (45). We believe the lack of S-COMT activity of our inhibitors contributes to the lower in vivo potency in the rat when compared to tolcapone. In clinical usage, MB-COMT selectivity may actually be an advantage for our compounds. In humans, and nonhuman primates, roughly 85% of total brain COMT is the MB-COMT form (46). Moreover, given the enzyme kinetics and endogenous dopamine concentrations in the brain, MB-COMT is most likely responsible for more than 85% of total brain COMT activity in humans (11, 47).

A critical outstanding question concerning the utility of COMT inhibitors is which clinical indication is most likely to be improved by treatment. Small, short-duration pilot studies have suggested potential efficacy in multiple disorders (24, 25, 44). In contrast, other studies have shown that in certain circumstances COMT inhibition has no effect on cognitive function and can actually impair cognition (22, 23). However, there is a convergence of data suggesting that disorders characterized by impaired executive function and/or impulse control, including traumatic brain injury (TBI), behavioral variant frontotemporal dementia (bvFTD), and ADHD may be good candidates for a COMT inhibitor clinical trial. To be clear, we do not believe that a COMT inhibitor strategy would be disease-modifying in these indications. However, there are currently no treatments available to significantly improve the chronic symptoms of TBI or bvFTD and there is a still a need for effective ADHD treatments without stimulant effects and abuse liability (48-50). LIBD-1 has shown significant efficacy in our preclinical models to suggest it may be able to serve as the COMT inhibitor to test these hypotheses.

## Funding and Disclosure

This work was funded by US National Institutes of Health grant R01MH107126 and The Lieber Institute for Brain Development. IPB and JCB are inventors on patents that include the novel COMT inhibitors (WO2016123576 and WO2017091818). HLR, RSK, LP, and SCC are employees of RenaSci Ltd. The remaining authors have nothing to disclose.

## Acknowledgments

The authors thank Michael Poslusney and Noelle White for technical assistance, Richard Brammer for statistical analysis of the microdialysis data, Elizabeth Tunbridge for advice on the ASST assay, and Daniel R. Weinberger for constructive comments on a draft of this manuscript.

