## Supplemental Information for "Novel, non-nitrocatechol COMT inhibitors modulate dopamine neurotransmission in the frontal cortex and improve cognitive flexibility"

### Supplemental Methods

*HPLC analysis of microdialysis samples*

Detection and subsequent quantification of DA, DOPAC and HVA in the dialysis samples was achieved by reverse-phase, ion-pair HPLC coupled with electrochemical detection and involved the use of an ALEXYS monoamine analyser (Antec Scientific, Zoeterwoude, The Netherlands). The system consisted of two separate analytical columns (ALF-115, 150 mm × 1 mm internal diameter) that shared a dual-loop autosampler, allowing for one sample to be simultaneously analysed by two systems optimized for different neurotransmitters. One column separated DOPAC and HVA, while the other separated DA. Two solvent delivery pumps (LC110) were used to circulate the respective mobile phases (DOPAC and HVA: 50 mM citric acid, 50 mM phosphoric acid, 8 mM NaCl, 0.1 mM EDTA, 6.9 mM 1-octanesulfonic acid, 10% methanol, pH 3.25; DA: 50 mM phosphoric acid, 8 mM NaCl, 0.1 mM EDTA, 10.4 mM 1-octane sulphonic acid, 20% methanol, pH 6.0) at a flow rate of 50 µL/min. An Antec in-line degassing unit was used to remove air. Samples (10 µl) were injected onto the columns via an autosampler (AS 110) with a cooling tray set at 4°C. Antec DECADE II electrochemical detectors were used comprising Antec micro VT03 cells employing a high density, glassy carbon working electrode (+0.59 V for DOPAC and HVA; +0.3 V for DA) combined with an Antec ISAAC reference electrode. The electrode signal was integrated using Antec's CLARITY data acquisition system. Individual stock solutions of DA, DOPAC and HVA (1.0 mM) were prepared by their dissolution in a mixture of equal quantities of deionized water and 0.1 M perchloric acid and stored at 4°C. A working solution containing all the transmitters and metabolites was prepared daily by dilution in aCSF.

*Probe location verification*

At the end of the experiment rats were euthanized and their brains were rapidly removed and stored in a 10% v/v formalin saline solution for a minimum of 5 days. Brain sections were cut using a vibratome and probe placements were localized with reference to a stereotaxic atlas (Paxinos and Watson, 1986)

*Attentional set-shifting task*

*Apparatus*

The apparatus is a modified version of an arena previous described (33). Briefly, the apparatus was a customized clear Plexiglas arena (70 x 40 x 18 cm (L x W x H); K and J Fabrications, Aberdeen, MD, USA) with an opaque Plexiglas panel dividing one third of the cage into two equally sized choice areas. The choice areas are separated from the start box via a removable opaque Plexiglas panel. Water was provided for rats in the start box.

*Odor/Digging exemplars*

Rats dug in porcelain, 3-ounce ramekin bowls (Norpro, Everett, WA, USA). Blotting paper (Maiko, Kyoto, Japan) was attached to the inner rim of the bowls with adhesive tape. The rewards used in this protocol were fresh Honey Nut Cheerio® halves (General Mills, Minneapolis, MN, USA). Cheerios® were stored in their original bag and held in resealable, plastic storage containers. The supply of Cheerios® was replaced every two weeks as digging behavior greatly decreased when the Cheerio® rewards became stale. Scented oils (Bakto Flavors, North Brunswick, NJ, USA) were applied to the blotting paper. Digging media consisted of small materials used to cover rewards in the bowls (Supplemental Table 1). Bowls were filled with the designated medium and blotting paper was affixed to each bowl's interior rim prior to the beginning of each attentional set-shifting session, minimizing inter-stage breaks as much as possible. Following the conclusion of any habituation, training or testing, blotting paper was removed and bowls were cleaned thoroughly.

*Pre-habituation*

Rats were given a bowl of corn cob bedding with three buried rewards in their home cage the day before habituation. This ensured that rats were familiar with digging for and eating rewards before testing began.

*Habituation (Day 1)*

Habituation was performed one week following the start of food restriction. Rats were allowed to acclimate to the testing room for one hour. An unscented bowl baited with three rewards was placed in the home cage and the rat was allowed to eat all rewards from the bowl.  Next, an unscented bowl with one reward was placed in the home cage. Following retrieval, the bowl was re-baited with a reward and with increasing amounts of digging medium (corn cob bedding). This process was repeated six times in the home cage allowing the rat to eat the reward each trial. These steps were repeated in the testing apparatus, beginning with a baited bowl with no digging media present in either choice area and ending with deeply buried rewards. After rats recovered buried rewards in each choice area they were ushered back into the starting box by gently rattling the removable divider. If a rat failed to complete habituation criterion in its first day, re-habituation was performed for up to two extra days.

*Training (Day 2)*

On the day following habituation, rats were trained on a series of two simple discriminations based on an odor cue (vanilla/banana) or a digging medium cue (black tea/sphagnum moss) to a criterion of six consecutive correct trials. The digging medium for the odor discrimination training consisted of home cage corn cob bedding.  The bowl marked with the 'positive' cue (i.e. vanilla$_{+}$, black tea$_{+}$) always contained a reward, while the bowl marked with the other cue (banana$_{-}$, sphagnum moss$_{-}$) never contained a reward. The same discriminations were presented to rats in the same order and were not used again during testing.  The negative bowl was masked by the addition of a small amount of fresh Honey Nut Cheerio® powder before each third trial. Placement of positive and negative bowls was balanced across the two choice areas during training. Digging media and odor cues are listed in Supplemental Table 1.

A trial was initiated by raising the starting divider giving the rat access to both choice areas. The first four trials in each discrimination stage were discovery trials where rats were allowed to dig in both choice areas without recording response accuracy. Discovery trials ended when they consumed the reward and explored the unbaited bowl. Trials continued beyond the discovery trials until a criterion of six consecutive correct trials was reached. Latency to dig and response accuracy were recorded for each trial following the discovery phase. An incorrect dig signaled the end of the trial- the rewarded bowl was removed and rats were ushered back to the starting box. Rats were allowed to climb over and investigate the bowls; a dig was only scored if the rat broke the surface by purposefully displacing digging medium with either its paws or nose. Occasionally, rats would mark the bowls with urine, possibly in attempts to use a mediating behavior to identify reward- containing bowls. In this situation, digging media for both bowls were discarded and replaced with fresh media before continuing the session. Once again, if a rat failed to complete training in its first day, re-training was performed for up to two extra days before that rat was removed from the study.

*Testing (Day 3)*

Rats performed a single attentional set shifting test the day following training. Testing was conducted in the same manner as training. Briefly, trials were initiated by the raising of the divider, the first four trials being unscored discovery trials, and a criterion of six consecutive correct responses was necessary to pass a stage. Rats were dosed with vehicle, tolcapone, LIBD-1, or LIBD-3 and returned to their home cage for one hour until testing.

Rats performed a series of seven discriminations in the following order: simple discrimination (SD), compound discrimination (CD), compound reversal (CDR), intradimensional discrimination (ID), intradimensional reversal (IDR), extradimensional discrimination (ED) and extradimensional reversal (EDR) (Table 1). The simple discrimination (SD) consisted of bowls differing by one of two dimensions (odor or digging medium). The compound discrimination stage (CD) introduced the second irrelevant dimension while the correct and incorrect exemplars were the same. The relevant dimension remained consistent during the compound dimension reversal stage (CDR) but the correct and incorrect exemplars were swapped so that the previously rewarded exemplar was no longer rewarded and vice versa. Following completion of the CDR the relevant dimension remained consistent in the intradimensional discrimination stage (ID) while a novel sets of odor and digging medium exemplars were used. As with the CDR stage, in the IDR stage the rewarded and non-rewarded exemplars were swapped. The previously relevant and irrelevant dimensions were swapped during the extradimensional discrimination (ED) with novel sets of odor and digging medium exemplars. Finally, the EDR stage, where correct and incorrect exemplars were swapped, was initiated following the completion of the ED. Once a rat reached a criterion of 6 correct trials in a row during the EDR the task was terminated and rats were returned to their home cages.

### Supplemental Table 1

| Phase | Odors | Media | Media Suppliers |
|---|---|---|---|
| Training | **Vanilla**/Banana | **Black Tea**/Long Fibered Sphagnum Moss | Positively Tea Company, Sellinsgrove, PA, USA/Mosser Lee Company, Millston, WI, USA |
| SD | **Rum**/Irish Crème | Home Bedding (corn cob) | Shepherd Specialty Papers, Milford, NJ, USA |
| CD | **Rum**/Irish Crème | Living World® Pine Shavings/Carefresh® Animal Bedding | Rolf C. Hagen, Mansfield, MA, USA/Healthy Pet, Ferndale, WA, USA |
| CDR | Rum/**Irish Crème** | Living World® Pine Shavings/Carefresh® Animal Bedding | Rolf C. Hagen, Mansfield, MA, USA/Healthy Pet, Ferndale, WA, USA |
| ID | **Almond**/Lemon | Blue Shredded Confetti/Pure Comfort® Animal Bedding | Amscan/Oxbow Animal Health, Omaha, NE, USA |
| IDR | Almond/**Lemon** | Blue Shredded Confetti/Pure Comfort® Animal Bedding | Amscan/Oxbow Animal Health, Omaha, NE, USA |
| ED | Orange/Anise | **Red Gravel**/White Gravel | PetSmart, Phoenix, AZ, USA |
| EDR | Orange/Anise | Red Gravel/**White Gravel** | PetSmart, Phoenix, AZ, USA |

**Bold** represents correct exemplar for each stage

### Supplemental Table 2 Dopamine results in the prefrontal cortex (fmol/5 µl)

| Time period | Treatment | n | Mean | SEM | % of baseline | | Comparison vs vehicle | |
|---|---|---|---|---|---|---|---|---|
| | | | | | Mean | SE | % change | p |
| -80 to -60 mins | Vehicle 10 ml/kg po | 8 | 1.02 | 0.06 | 100.7 | 14.1 | . | |
| | Tolcapone 15 mg/kg ip | 8 | 0.96 | 0.06 | 95.2 | 13.8 | -5.5 | 0.664 |
| | LIBD-1 100 mg/kg po | 8 | 0.99 | 0.12 | 97.9 | 17.3 | -2.8 | 0.827 |
| | LIBD-3 100 mg/kg po | 8 | 1.10 | 0.10 | 108.8 | 17.4 | 8.0 | 0.568 |
| -60 to -40 mins | Vehicle 10 ml/kg po | 8 | 0.99 | 0.06 | 98.1 | 14.1 | . | |
| | Tolcapone 15 mg/kg ip | 7 | 1.17 | 0.10 | 116.2 | 17.6 | 18.4 | 0.178 |
| | LIBD-1 100 mg/kg po | 8 | 1.07 | 0.11 | 106.1 | 17.1 | 8.1 | 0.514 |
| | LIBD-3 100 mg/kg po | 8 | 1.00 | 0.08 | 98.5 | 14.8 | 0.4 | 0.977 |
| -40 to -20 mins | Vehicle 10 ml/kg po | 7 | 0.99 | 0.04 | 97.5 | 13.2 | . | |
| | Tolcapone 15 mg/kg ip | 8 | 0.90 | 0.04 | 89.0 | 12.2 | -8.6 | 0.373 |
| | LIBD-1 100 mg/kg po | 8 | 0.94 | 0.07 | 92.9 | 13.8 | -4.6 | 0.639 |
| | LIBD-3 100 mg/kg po | 8 | 0.91 | 0.08 | 90.2 | 14.0 | -7.5 | 0.452 |
| -20 to 0 mins | Vehicle 10 ml/kg po | 8 | 1.13 | 0.09 | 111.6 | 16.9 | . | |
| | Tolcapone 15 mg/kg ip | 8 | 0.88 | 0.10 | 86.6 | 15.2 | -22.4 | 0.181 |
| | LIBD-1 100 mg/kg po | 8 | 1.06 | 0.12 | 105.0 | 18.1 | -5.9 | 0.746 |
| | LIBD-3 100 mg/kg po | 8 | 0.87 | 0.15 | 85.7 | 18.4 | -23.1 | 0.176 |
| 0 to 20 mins | Vehicle 10 ml/kg po | 8 | 1.27 | 0.06 | 125.8 | 17.1 | . | |
| | Tolcapone 15 mg/kg ip | 8 | 1.26 | 0.10 | 124.2 | 19.0 | -1.3 | 0.916 |
| | LIBD-1 100 mg/kg po | 8 | 1.15 | 0.07 | 113.6 | 16.1 | -9.7 | 0.405 |
| | LIBD-3 100 mg/kg po | 8 | 1.22 | 0.15 | 120.6 | 21.1 | -4.1 | 0.734 |
| 20 to 40 mins | Vehicle 10 ml/kg po | 8 | 1.11 | 0.13 | 109.6 | 18.9 | . | |
| | Tolcapone 15 mg/kg ip | 7 | 1.35 | 0.27 | 133.2 | 31.5 | 21.6 | 0.545 |
| | LIBD-1 100 mg/kg po | 8 | 1.04 | 0.12 | 102.7 | 17.9 | -6.3 | 0.834 |
| | LIBD-3 100 mg/kg po | 8 | 0.79 | 0.26 | 78.2 | 28.0 | -28.7 | 0.293 |
| 40 to 60 mins | Vehicle 10 ml/kg po | 8 | 1.01 | 0.07 | 99.7 | 14.3 | . | |
| | Tolcapone 15 mg/kg ip | 8 | 0.86 | 0.16 | 85.0 | 18.8 | -14.7 | 0.549 |
| | LIBD-1 100 mg/kg po | 8 | 1.03 | 0.27 | 101.7 | 30.0 | 2.1 | 0.939 |
| | LIBD-3 100 mg/kg po | 8 | 1.08 | 0.16 | 106.7 | 20.7 | 7.0 | 0.803 |
| 60 to 80 mins | Vehicle 10 ml/kg po | 7 | 1.17 | 0.09 | 116.0 | 17.2 | . | |
| | Tolcapone 15 mg/kg ip | 8 | 1.10 | 0.13 | 108.9 | 19.0 | -6.2 | 0.725 |
| | LIBD-1 100 mg/kg po | 8 | 1.01 | 0.09 | 100.0 | 15.7 | -13.8 | 0.416 |
| | LIBD-3 100 mg/kg po | 8 | 0.97 | 0.16 | 95.9 | 19.9 | -17.3 | 0.311 |
| 80 to 100 mins | Vehicle 10 ml/kg po | 8 | 1.03 | 0.11 | 102.2 | 16.8 | . | |
| | Tolcapone 15 mg/kg ip | 8 | 0.99 | 0.18 | 97.8 | 21.9 | -4.4 | 0.843 |
| | LIBD-1 100 mg/kg po | 7 | 1.09 | 0.22 | 107.8 | 25.8 | 5.5 | 0.819 |
| | LIBD-3 100 mg/kg po | 8 | 1.03 | 0.12 | 101.3 | 17.6 | -0.9 | 0.970 |
| 100 to 120 mins | Vehicle 10 ml/kg po | 8 | 1.08 | 0.13 | 106.3 | 18.6 | . | |
| | Tolcapone 15 mg/kg ip | 8 | 0.98 | 0.12 | 96.5 | 17.2 | -9.2 | 0.705 |
| | LIBD-1 100 mg/kg po | 8 | 0.93 | 0.16 | 92.2 | 19.8 | -13.3 | 0.581 |
| | LIBD-3 100 mg/kg po | 8 | 0.98 | 0.24 | 97.1 | 27.0 | -8.6 | 0.732 |

Means are adjusted for differences between treatment groups at baseline. SEM's are calculated from the residuals of the statistical model. % of baseline calculated from the adjusted means.
P values are for the comparison to vehicle by the multiple t test.

**Supplemental Table 3  DOPAC results in the prefrontal cortex (fmol/5 µl)**

| Time period | Treatment | n | Mean | SEM | % of baseline | | Comparison vs vehicle | |
|---|---|---|---|---|---|---|---|---|
| | | | | | Mean | SE | % change | p |
| -80 to -60 mins | Vehicle 10 ml/kg po | 8 | 110 | 1 | 101.9 | 11.7 | . | |
| | Tolcapone 15 mg/kg ip | 7 | 107 | 2 | 99.4 | 11.4 | -2.5 | 0.450 |
| | LIBD-1 100 mg/kg po | 8 | 109 | 2 | 100.9 | 11.7 | -0.9 | 0.769 |
| | LIBD-3 100 mg/kg po | 8 | 108 | 4 | 100.2 | 11.9 | -1.7 | 0.630 |
| -60 to -40 mins | Vehicle 10 ml/kg po | 8 | 107 | 2 | 98.9 | 11.3 | . | |
| | Tolcapone 15 mg/kg ip | 7 | 103 | 6 | 95.3 | 12.0 | -3.6 | 0.553 |
| | LIBD-1 100 mg/kg po | 8 | 110 | 2 | 102.1 | 11.8 | 3.3 | 0.596 |
| | LIBD-3 100 mg/kg po | 8 | 103 | 6 | 95.8 | 12.3 | -3.1 | 0.639 |
| -40 to -20 mins | Vehicle 10 ml/kg po | 7 | 102 | 2 | 94.2 | 10.9 | . | |
| | Tolcapone 15 mg/kg ip | 7 | 108 | 6 | 100.1 | 12.7 | 6.3 | 0.311 |
| | LIBD-1 100 mg/kg po | 8 | 99 | 2 | 91.9 | 10.6 | -2.4 | 0.675 |
| | LIBD-3 100 mg/kg po | 8 | 107 | 5 | 98.8 | 12.2 | 4.9 | 0.442 |
| -20 to 0 mins | Vehicle 10 ml/kg po | 8 | 114 | 4 | 105.8 | 12.6 | . | |
| | Tolcapone 15 mg/kg ip | 7 | 109 | 4 | 100.6 | 12.1 | -5.0 | 0.504 |
| | LIBD-1 100 mg/kg po | 8 | 103 | 5 | 95.6 | 11.9 | -9.7 | 0.175 |
| | LIBD-3 100 mg/kg po | 8 | 109 | 7 | 101.0 | 13.3 | -4.6 | 0.563 |
| 0 to 20 mins | Vehicle 10 ml/kg po | 8 | 118 | 3 | 109.1 | 12.7 | . | |
| | Tolcapone 15 mg/kg ip | 7 | 115 | 8 | 106.7 | 14.2 | -2.3 | 0.796 |
| | LIBD-1 100 mg/kg po | 8 | 107 | 8 | 99.4 | 13.6 | -8.9 | 0.286 |
| | LIBD-3 100 mg/kg po | 8 | 106 | 6 | 98.1 | 12.4 | -10.1 | 0.267 |
| 20 to 40 mins | Vehicle 10 ml/kg po | 8 | 114 | 4 | 105.5 | 12.6 | . | |
| | Tolcapone 15 mg/kg ip | 6 | 159 | 7 | 147.2 | 18.1 | 39.4 | <0.001*** |
| | LIBD-1 100 mg/kg po | 8 | 126 | 9 | 117.1 | 15.5 | 11.0 | 0.186 |
| | LIBD-3 100 mg/kg po | 7 | 139 | 8 | 129.1 | 16.5 | 22.3 | 0.026* |
| 40 to 60 mins | Vehicle 10 ml/kg po | 8 | 112 | 2 | 104.0 | 11.9 | . | |
| | Tolcapone 15 mg/kg ip | 7 | 233 | 6 | 215.7 | 25.1 | 107.4 | <0.001*** |
| | LIBD-1 100 mg/kg po | 8 | 168 | 17 | 155.5 | 23.5 | 49.4 | <0.001*** |
| | LIBD-3 100 mg/kg po | 7 | 194 | 16 | 180.1 | 25.4 | 73.1 | <0.001*** |
| 60 to 80 mins | Vehicle 10 ml/kg po | 7 | 101 | 6 | 93.9 | 12.1 | . | |
| | Tolcapone 15 mg/kg ip | 7 | 237 | 33 | 219.4 | 39.3 | 133.7 | <0.001*** |
| | LIBD-1 100 mg/kg po | 8 | 207 | 11 | 191.6 | 24.1 | 104.1 | <0.001*** |
| | LIBD-3 100 mg/kg po | 8 | 184 | 34 | 170.1 | 36.9 | 81.1 | 0.008** |
| 80 to 100 mins | Vehicle 10 ml/kg po | 8 | 103 | 4 | 95.1 | 11.5 | . | |
| | Tolcapone 15 mg/kg ip | 6 | 309 | 10 | 285.9 | 33.8 | 200.8 | <0.001*** |
| | LIBD-1 100 mg/kg po | 7 | 216 | 10 | 199.7 | 24.5 | 110.1 | <0.001*** |
| | LIBD-3 100 mg/kg po | 8 | 221 | 13 | 204.3 | 26.2 | 114.9 | <0.001*** |
| 100 to 120 mins | Vehicle 10 ml/kg po | 8 | 105 | 3 | 97.0 | 11.3 | . | |
| | Tolcapone 15 mg/kg ip | 6 | 322 | 12 | 297.7 | 35.6 | 206.8 | <0.001*** |
| | LIBD-1 100 mg/kg po | 8 | 250 | 10 | 231.1 | 27.8 | 138.1 | <0.001*** |
| | LIBD-3 100 mg/kg po | 8 | 218 | 26 | 202.0 | 33.2 | 108.1 | <0.001*** |

Means are adjusted for differences between treatment groups at baseline. SEM's are calculated from the residuals of the statistical model. % of baseline calculated from the adjusted means.
P values are for the comparison to vehicle by the multiple t test. *p<0.05, **p<0.01, ***p<0.001

**Supplemental Table 4  HVA results in the prefrontal cortex (fmol/5 µl)**

| Time period | Treatment | n | Mean | SEM | % of baseline | | Comparison vs vehicle | |
|---|---|---|---|---|---|---|---|---|
| | | | | | Mean | SE | % change | p |
| -80 to -60 mins | Vehicle 10 ml/kg po | 8 | 281 | 4 | 103.5 | 9.9 | . | |
| | Tolcapone 15 mg/kg ip | 7 | 278 | 4 | 102.2 | 9.8 | -1.2 | 0.724 |
| | LIBD-1 100 mg/kg po | 8 | 279 | 6 | 102.7 | 10.0 | -0.8 | 0.815 |
| | LIBD-3 100 mg/kg po | 7 | 292 | 11 | 107.3 | 11.0 | 3.7 | 0.349 |
| -60 to -40 mins | Vehicle 10 ml/kg po | 8 | 274 | 3 | 100.8 | 9.7 | . | |
| | Tolcapone 15 mg/kg ip | 7 | 261 | 11 | 96.0 | 9.9 | -4.8 | 0.396 |
| | LIBD-1 100 mg/kg po | 8 | 283 | 4 | 104.3 | 10.0 | 3.4 | 0.551 |
| | LIBD-3 100 mg/kg po | 8 | 259 | 16 | 95.1 | 10.8 | -5.7 | 0.353 |
| -40 to -20 mins | Vehicle 10 ml/kg po | 7 | 255 | 4 | 94.0 | 9.0 | . | |
| | Tolcapone 15 mg/kg ip | 7 | 271 | 14 | 99.6 | 10.7 | 5.9 | 0.354 |
| | LIBD-1 100 mg/kg po | 8 | 248 | 5 | 91.2 | 8.9 | -2.9 | 0.618 |
| | LIBD-3 100 mg/kg po | 8 | 262 | 15 | 96.5 | 10.7 | 2.7 | 0.686 |
| -20 to 0 mins | Vehicle 10 ml/kg po | 8 | 296 | 12 | 108.7 | 11.3 | . | |
| | Tolcapone 15 mg/kg ip | 7 | 270 | 12 | 99.4 | 10.5 | -8.6 | 0.234 |
| | LIBD-1 100 mg/kg po | 8 | 268 | 9 | 98.5 | 10.0 | -9.4 | 0.179 |
| | LIBD-3 100 mg/kg po | 8 | 275 | 19 | 101.3 | 11.8 | -6.8 | 0.384 |
| 0 to 20 mins | Vehicle 10 ml/kg po | 8 | 299 | 12 | 110.0 | 11.3 | . | |
| | Tolcapone 15 mg/kg ip | 7 | 282 | 19 | 103.8 | 12.0 | -5.7 | 0.482 |
| | LIBD-1 100 mg/kg po | 8 | 283 | 15 | 104.0 | 11.4 | -5.5 | 0.480 |
| | LIBD-3 100 mg/kg po | 8 | 258 | 15 | 95.0 | 10.6 | -13.7 | 0.109 |
| 20 to 40 mins | Vehicle 10 ml/kg po | 8 | 290 | 9 | 106.8 | 10.7 | . | |
| | Tolcapone 15 mg/kg ip | 6 | 233 | 10 | 85.6 | 8.9 | -19.8 | 0.006** |
| | LIBD-1 100 mg/kg po | 8 | 270 | 15 | 99.5 | 10.9 | -6.8 | 0.303 |
| | LIBD-3 100 mg/kg po | 7 | 232 | 14 | 85.3 | 9.5 | -20.1 | 0.007** |
| 40 to 60 mins | Vehicle 10 ml/kg po | 8 | 294 | 9 | 108.2 | 10.8 | . | |
| | Tolcapone 15 mg/kg ip | 7 | 164 | 8 | 60.3 | 6.4 | -44.3 | <0.001*** |
| | LIBD-1 100 mg/kg po | 8 | 256 | 18 | 94.2 | 11.2 | -12.9 | 0.090 |
| | LIBD-3 100 mg/kg po | 7 | 172 | 11 | 63.3 | 7.1 | -41.5 | <0.001*** |
| 60 to 80 mins | Vehicle 10 ml/kg po | 8 | 276 | 15 | 101.6 | 11.2 | . | |
| | Tolcapone 15 mg/kg ip | 7 | 85 | 12 | 31.1 | 5.4 | -69.4 | <0.001*** |
| | LIBD-1 100 mg/kg po | 8 | 203 | 14 | 74.7 | 8.9 | -26.5 | 0.181 |
| | LIBD-3 100 mg/kg po | 8 | 99 | 26 | 36.6 | 10.0 | -64.0 | <0.001*** |
| 80 to 100 mins | Vehicle 10 ml/kg po | 8 | 281 | 12 | 103.4 | 10.7 | . | |
| | Tolcapone 15 mg/kg ip | 6 | 61 | 9 | 22.6 | 3.9 | -78.1 | <0.001*** |
| | LIBD-1 100 mg/kg po | 7 | 161 | 11 | 59.1 | 7.0 | -42.9 | 0.003** |
| | LIBD-3 100 mg/kg po | 8 | 73 | 12 | 27.0 | 5.1 | -73.9 | <0.001*** |
| 100 to 120 mins | Vehicle 10 ml/kg po | 8 | 264 | 16 | 97.3 | 10.9 | . | |
| | Tolcapone 15 mg/kg ip | 6 | 41 | 6 | 15.1 | 2.6 | -84.4 | <0.001*** |
| | LIBD-1 100 mg/kg po | 8 | 152 | 16 | 55.8 | 8.0 | -42.6 | 0.057 |
| | LIBD-3 100 mg/kg po | 8 | 44 | 15 | 16.3 | 5.6 | -83.2 | <0.001*** |

Means are adjusted for differences between treatment groups at baseline. SEM's are calculated from the residuals of the statistical model. % of baseline calculated from the adjusted means.
P values are for the comparison to vehicle by the multiple t test. *p<0.05, **p<0.01, ***p<0.001

### Supplemental Table 5 – Extreme values (|z|>4)

| Measurement | Rat | Time (mins) | Value | z |
|---|---|---|---|---|
| Dopamine | 32A | 80 | 0.07 | -7.29 |
| Dopamine | 22D | 40 | 0.03 | -5.78 |
| DOPAC | 2C | 20 | 44 | -4.35 |
| DOPAC | 2C | 60 | 58 | -5.28 |
| DOPAC | 20D | -40 | 27 | -4.88 |
| DOPAC | 20D | 0 | 74 | 4.44 |
| DOPAC | 20D | 80 | 20 | -5.76 |
| DOPAC | 20D | 120 | 42 | -6.14 |
| HVA | 20D | -40 | 76 | -6.44 |
| HVA | 20D | 0 | 212 | 4.46 |
| HVA | 20D | 80 | 8 | -7.75 |
| HVA | 20D | 100 | 12 | -5.02 |
| HVA | 20D | 120 | 3 | -5.29 |

z is the 'studentised residual' from the statistical model - for normally distributed data it is less than -4 or greater than 4 for about 1 in 10,000 observations.
